# Extracellular ATP released from *Candida albicans* activates non-peptidergic neurons to augment host defense

**DOI:** 10.1101/2020.01.27.921049

**Authors:** Tara N Edwards, Shiqun Zhang, Andrew Liu, Jonathan A. Cohen, Paul Yifan Zhou, Selene Mogavero, Bernhard Hube, Judith Berman, Marie-Elisabeth Bougnoux, Alicia R. Mathers, Sarah L. Gaffen, Kathryn M. Albers, H. Richard Koerber, Brian M. Davis, Christophe D’Enfert, Daniel H Kaplan

## Abstract

Intestinal microbes release ATP to modulate local immune responses. Herein we demonstrates that *Candida albicans*, an opportunistic commensal fungus, also modulates immune responses via secretion of ATP. We found that ATP secretion from *C. albicans* varied between standard laboratory strains. A survey of eighty-nine clinical isolates revealed heterogeneity in ATP secretion, independent of growth kinetics and intracellular ATP levels. Isolates from blood released less ATP than commensals, suggesting that ATP secretion assists with commensalism. To confirm this, cohorts of mice were infected with strains matched for origin, and intracellular ATP concentration, but high or low extracellular ATP. In all cases fungal burden was inversely correlated with ATP secretion. Mice lacking P2RX7, the key ATP receptor expressed by immune cells in the skin, showed no alteration in fungal burden. Rather, treatments with a P2RX2/3 antagonist result in increased fungal burden. P2RX2/3 is expressed by non-peptidergic neurons that terminate in the epidermis. Cultured sensory neurons flux Ca^2+^ when exposed to supernatant from heat-killed *C. albicans* (HKCA), and these non-peptidergic fibers are the dominant subset that respond to HKCA. Ca^2+^ flux, but not CGRP-release, can be abrogated by pretreatment of HKCA supernatant with apyrase. To determine whether non-peptidergic neurons participate in host defense, we generated MRGPRD-DTR mice. Infection in these mice resulted in increased CFU only for those *C. albicans* strains with high ATP secretion. Taken together, our findings indicate that *C. albicans* releases ATP, which is recognized by non-peptidergic nerves in the skin resulting in augmented anti-*Candida* immune responses.

**Author Summary:** Bacterial release of ATP has been shown to modulate immune responses. *Candida albicans* displays heterogeneity in ATP release among laboratory strains and commensal clinical isolates release more ATP than invasive isolates. *C. albicans* strains with high ATP secretion show lower fungal burden following epicutaneous infection. Mice lacking P2RX7, the key ATP receptor expressed by immune cells, showed no alteration in fungal burden. In contrast, treatment with P2RX2/3 antagonists resulted in increased fungal burden. P2RX3 is expressed by a subset of non-peptidergic neurons that terminate in the epidermis. These non-peptidergic fibers are the predominant responders when cultured sensory neurons are exposed to heat-killed *C. albicans in vitro*. Mice lacking non-peptidergic neurons have increased infection when exposed to high but not low ATP-secreting isolates of *C. albicans.* Taken together, our findings indicate that *C. albicans* releases ATP which is recognized by non-peptidergic nerves in the skin resulting in augmented anti-*Candida* immune responses.

**Bullet points:**

- ATP released from heat killed *C. albicans* activates non-peptidergic sensory neurons
- Live *C. albicans* clinical isolates release variable amounts of ATP
- Elevated levels of ATP released by *C. albicans* correlates with reduced infectivity *in vivo*
- MRGPRD-expressing cutaneous neurons are required for defense against ATP-secreting *C. albicans*

## Introduction

Skin is a barrier tissue that is exposed to both commensal microorganisms and pathogens. *Candida albicans* is a dimorphic fungus that typically grows as a commensal at barrier surfaces, but it can also become pathogenic. Pathenogenic *C. albicans* infection can manifest as chronic mucocutaneous candidiasis or disseminated candidiasis in immunocompromised individuals, leading to fungal sepsis in severe cases [1]. In healthy individuals, type-17 antifungal immunity limits *C. albicans* growth at barrier tissues [2]. Patients with genetic mutations in this pathway can suffer chronic mucocutaneous candidiasis [3]. In most mice strains, *C. albicans* is a foreign pathogen that is commonly used as a model to interrogate mechanisms of fungal immunity and host defense. As in humans, type-17 immunity mediates anti-candida responses at barrier tissues [2,4,5].

Immune recognition of *C. albicans* occurs through well described mechanisms involving mannose (mannose receptor [5]), mannoproteins (Toll-like receptor 4), phospholipomannan (Toll-like receptor 2 [6]), and β-mannosides (galectin-3 [7]). Of particular importance are β(1,3)-glucans that are detected by Dectin-1, a C-type lectin receptor on the surface of myeloid cells [8]. Together, recognition of these pathogen-associated molecular patterns (PAMPs) ultimately give rise to protective type-17 immunity [4].

Neurons of the somatosensory nervous system, particularly unmyelinated free nerve endings that project extensively throughout barrier tissues, are ideally located to detect danger signals. Unmyelinated C-fibers can be divided into multiple categories based on unique gene expression patterns [9,10]. Broadly speaking, these can be divided into peptidergic neurons that express TRPV1 and CGRP, and non-peptidergic neurons, many of which express the ATP receptor P2RX3 [11,12]. In skin, TRPV1-expressing neurons terminate primarily within the dermis and in the stratum spinosum, while terminals of nonpeptidergic neurons extend into the stratum granulosum [12]. During epicutaneous *C. albicans* infection, activation of TRPV1-expressing neurons is both necessary and sufficient to trigger the development of protective type-17 host defense through a mechanism that depends on CGRP [13,14].

Neurons directly recognize both bacteria and *C. albicans*. S*taphylococcus aureus* triggers neuron activation as measured by Ca^2+^ flux in sensory neurons [15]. *Streptococcus pyogenes* activates TRPV1^+^ neurons and triggers CGRP release [16]. Cutaneous neurons also detect *C. albicans*, but the mechanisms involved are less clear. *C. albicans* is sufficient to induce Ca^2+^ flux in both TRPV1-expressing peptidergic neurons and non-TRPV1 expressing neurons [13]. Dectin-1 and TRP channels are required for optimal CGRP release [17] and exposure of cultured dorsal root ganglia (DRG) neurons to soluble β-glucan is sufficient to evoke CGRP release [18]. Despite strong evidence that TRPV1-expressing neurons recognize pathogens and modulate host defense, little is known about the role of TRVP1-negative sensory neurons in host defense.

Most non-TRPV1 expressing neurons flux Ca^2+^ in response to *C. albicans* through an unknown mechanism [13]. Interestingly, ATP modulates β-glucan induced mechanical allodynia during *C. albicans* infection [18]. In this model, extracellular ATP (eATP) released from keratinocytes was suggested as the source. Notably, eATP is released by *E. coli* in the intestine as well as some other bacteria species and functions to modulate the development of local immune responses [19,20]. This raises the possibility that pathogen-derived eATP could trigger neuronal recognition of pathogens in skin.

Herein we show that C*. albicans* release eATP. The amount of eATP is highly variable between strains and positively correlates with the effectiveness of host defense that requires P2RX2/3 receptors. We also show that the major subset of non-peptidergic C-fiber neurons known to express P2RX3 are required for optimal host defense only against strains of *C. albicans* that express high levels of eATP. These data reveal an important unexpected host defense function for non-peptidergic neurons that relies on recognition of a novel *C. albicans*-derived PAMP.

## Results

### ATP from heat-killed C. albicans induces Ca^2+^ flux in cultured DRG neurons

In an effort to determine *C. albicans*-associated PAMPS responsible for activating cutaneous sensory neurons, we examined Ca^2+^ flux in Fura2-AM labeled cultured DRGs. *C. albicans* strain SC5314 was harvested at mid-log phase and washed into neuron recording buffer (NRB) prior to heat killing at 100° for 30-60 minutes. Heat killed *C. albicans* (HKCA) cells were separated into washed yeast cell bodies and effluence by centrifugation. Live neurons, identified by a response to 50mM potassium chloride (KCl), fluxed Ca^2+^ in response to HKCA effluence but not yeast cell bodies (**Fig 1A-B**). Approximately 45% of the 154 neurons examined responded to HKCA effluence. Ca^2+^ flux was evident within 1 second following application of effluence and responses were absent in Ca^2+^-free buffer (unpublished observation). Both capsaicin responsive (TRPV1+) and non-responsive (TRPV1-) neurons fluxed Ca^2+^ in the presence of HKCA effluence (**Fig 1A**). This is consistent with our prior observations and suggests that multiple populations of sensory afferent neurons respond to *C. albicans*.

**Fig 1:**
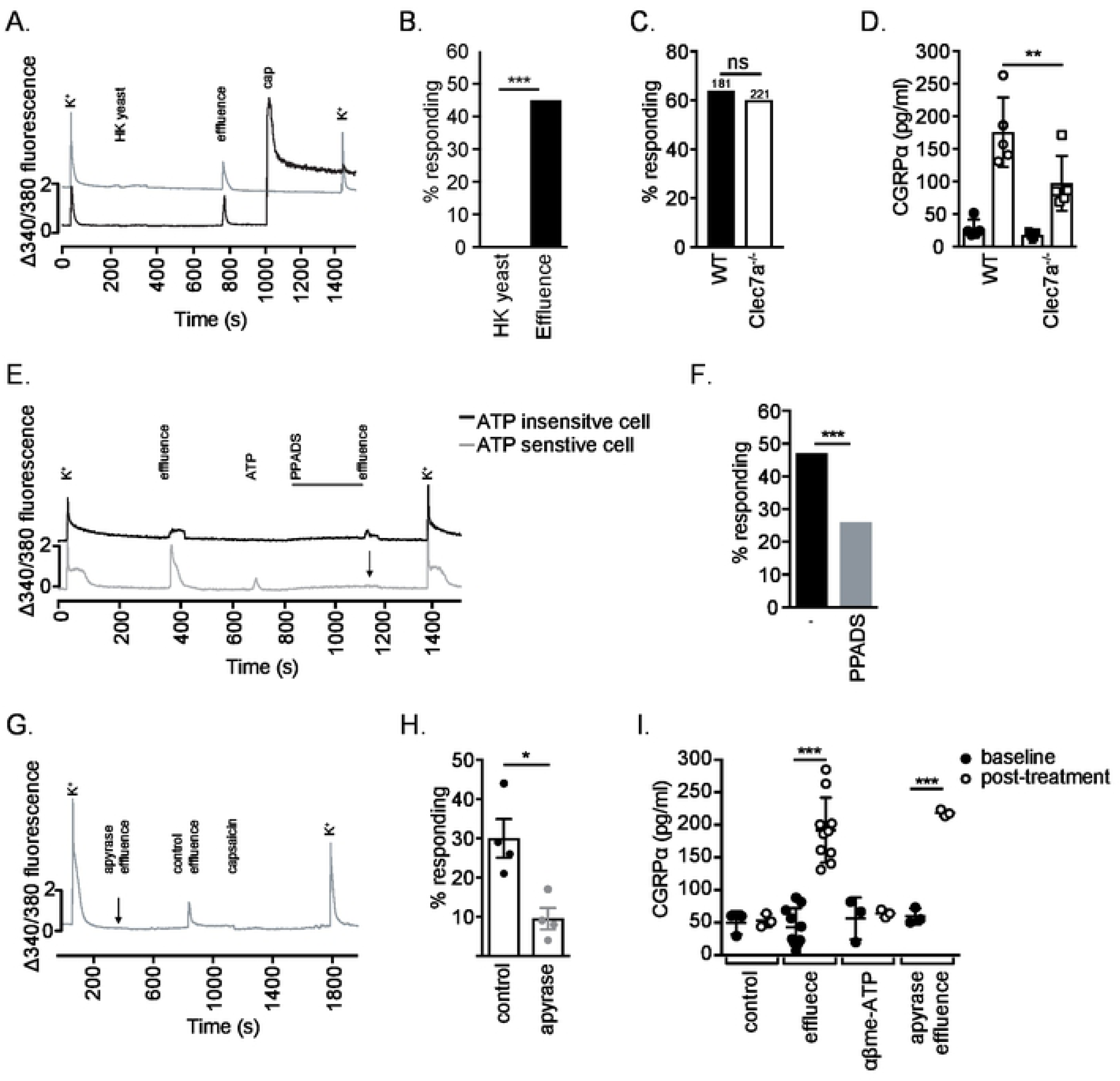
Sensory neurons detect *C. albicans* derived ATP. **(A)** Cultured DRG neurons were labeled with Fura2-AM and changes in intracellular Ca^2+^ in response to applied stimuli was measured as a change in 340/380 fluorescence over time. Example traces from two individual neurons indicate a change in intracellular Ca^2+^ upon treatment with effluence, but not HK SC5314 yeast, in both capsaicin-responsive and capsaicin non-responsive neurons. **(B)** Summary data as in (A) showing total numbers of lumbar DRG neurons responding to HKCA effluence but not to washed yeast. **(C)** Summary data of Ca^2+^ flux in DRG neurons isolated from *Clec7a*^−/−^ or WT mice exposed to HKCA effluence. **(D)** CGRP release from cultured DRG neurons isolated from WT and *Clec7a*^−/−^ mice following 10 minute exposure to HKCA effluence (open symbol) or immediately prior to HKCA exposure (closed symbols). **(E)** Ca^2+^ flux traces from two representative individual HKCA-responsive neurons, one ATP responsive (grey) and one ATP unresponsive (black). 10μM PPADS (P2XR2/3 inhibitor) was applied for 4 minutes (black bar) followed by challenge with HKCA effluence. **(F)** Summary data from (E) showing a 44% reduction in the number of neurons responding to HKCA effluence following PPADS incubation. **(G)** A representative Ca^2+^ trace from a single representative neuron that is unresponsive to apyrase treated HKCA effluence but responsive to control HKCA effluence. **(H)** Summary data from (G) showing a 68% reduction in the number of neurons responding to HKCA effluence treated with apyrase. **(I)** CGRP release from cultured DRG neurons isolated from WT mice obtained prior to stimulation (black circles) or 10 minutes following exposure to HKCA effluence, 100μM αβ-meATP, or apyrase treated HKCA effluence (open circles). Each symbol represents data from an individual well of cultured DRG (D, I) or pooled data from DRG isolated from an individual mouse (H). Significance was calculated using a chi-square test for Ca^2+^ flux assays (B, C, F) or Mann-Whitney (D, H, I). *p<0.05, \*\**p*<0.001, \*\*\**p*<0.0001. See also Fig S1.

β(1-3)-glucan is a well-defined component of the *C. albicans* yeast cell wall that drives recognition of *C. albicans* by cells of the innate immune system through interaction with the c-type lectin receptor Dectin-1 [21]. Although mRNA encoding for Dectin-1 (i.e. *Clec7a* mRNA) is not abundant in DRG single cell RNAseq datasets [9,22], incubation of cultured DRGs with *C. albicans*-derived soluble β-glucan is sufficient to trigger release of the neuropeptide CGRPα in a Dectin-1 dependent manner [18]. Unexpectedly, we observed that equal numbers of sensory neurons isolated from wild-type (WT) or *Clec7a*^−/−^ mice fluxed Ca^2+^ in response to HKCA effluence (**Fig 1C**). However, neurons isolated from *Clec7a*^−/−^ mice released significantly less CGRPα compared to neurons from WT mice (**Fig 1D**). These data support earlier reports that CGRPα release is at least partially dependent upon Dectin-1 [18], but also indicate that sensory neurons can be activated by a component in HKCA effluence other than β-glucan that is not dependant on Dectin-1.

The immediate Ca^2+^ flux observed in response to HKCA effluence (**Fig 1A**) suggests activation of a ligand gated ion channel. As many epidermal nonpeptidergic neurons express the P2RX3 receptor we examined the possible involvement of ATP [12]. To determine whether ATP could be responsible for Dectin-1 independent DRG activation, we first identified HKCA effluence-responsive cells based on Ca^2+^ flux (**Fig 1E**). Neurons were then tested for responsiveness to ATP by exposing them to 10μM ATP. PPADS, a selective P2 purinergic receptor antagonist that blocks activity of ligand-gated P2RX1-3 and P2RX5 receptors [23,24], was added to cultures prior to exposure to HKCA effluence. The presence of PPADS decreased the number total of neurons responding to HKCA effluence by 44% (**Fig 1E-F**). Notably, at least two kinds of HKCA effluence responsive neurons could be identified. Some neurons fluxed Ca^2+^ following exposure to ATP and their response to HKCA effluence could be inhibited by PPADS. Other neurons were unresponsive to ATP and their response to HKCA was not inhibited by PPADS.

To confirm that eATP in HKCA effluence was responsible for Ca^2+^ flux, we first determined that HKCA effluence contains approximately 10µM ATP and it could be efficiently reduced by treatment with apyrase, an ATP-diphosphohydrolase that catalyzes the hydrolysis of ATP to AMP (**S1A Fig**). Exposure of cultured DRGs to apyrase treated HKCA effluence reduced the number of responding neurons by 68% compared with HKCA effluence (**Figs 1G-H**). Notably, release of CGRPα was unaffected by treatment with apyrase (**Fig 1I**). Moreover, αβme-ATP, a selective P2RX1, P2RX3, and P2RX2/3 agonist [25], was not sufficient to elicit CGRPα release (**Fig 1I**). Taken together, these data support a model in which at least 2 subsets of neurons can respond to HKCA effluence. One releases CGRPα and is partially dependent on Dectin-1. This is likely the well-studied TRPV1^+^ expressing subset of cutaneous afferent neurons [13,14,18]. The other subset expresses a P2X receptor and is not linked to CGRPα release. These data also raise the exciting possibility that *C. albicans* secretes eATP, which could represent an unrecognized fungal PAMP that may participate in eliciting host defense.

### C. albicans strains secrete varying amounts of ATP

To determine whether live *C. albicans* releases eATP, mid-log-phase yeasts of *C. albicans* strain SC5314 were washed and suspended into PBS, and either incubated at 30°C or heat killed at 100°C for 1 hr. The quantity of eATP in the centrifuged supernatant of live *C. albicans* showed a significant amount of ATP that approached the level obtained by heat killing, thereby demonstrating that live *C. albicans* can release eATP (**Fig 2A**). To evaluate whether eATP resulted from cell death, we performed a kinetic analysis of eATP during growth in YPD broth. SC5314 yeast were inoculated into YPD and levels of eATP in the broth were analyzed over the course of a 12 hour 30° shaking incubation. eATP in the broth was first detectable after 2 hours and peaked at 6 hours during mid-log phase. The same samples were analyzed for cell viability by Propidium iodide staining and flow cytometry. Control HKCA were >99% PI+ while samples grown in YPD showed less than 0.5% staining at all time points tested (**Fig 2B, S2 Fig**). Thus, *C. albicans* SC5314 releases eATP during culture in both YPD and PBS that is unrelated to cell death, suggesting that eATP is actively secreted by viable cells.

**Fig 2:**
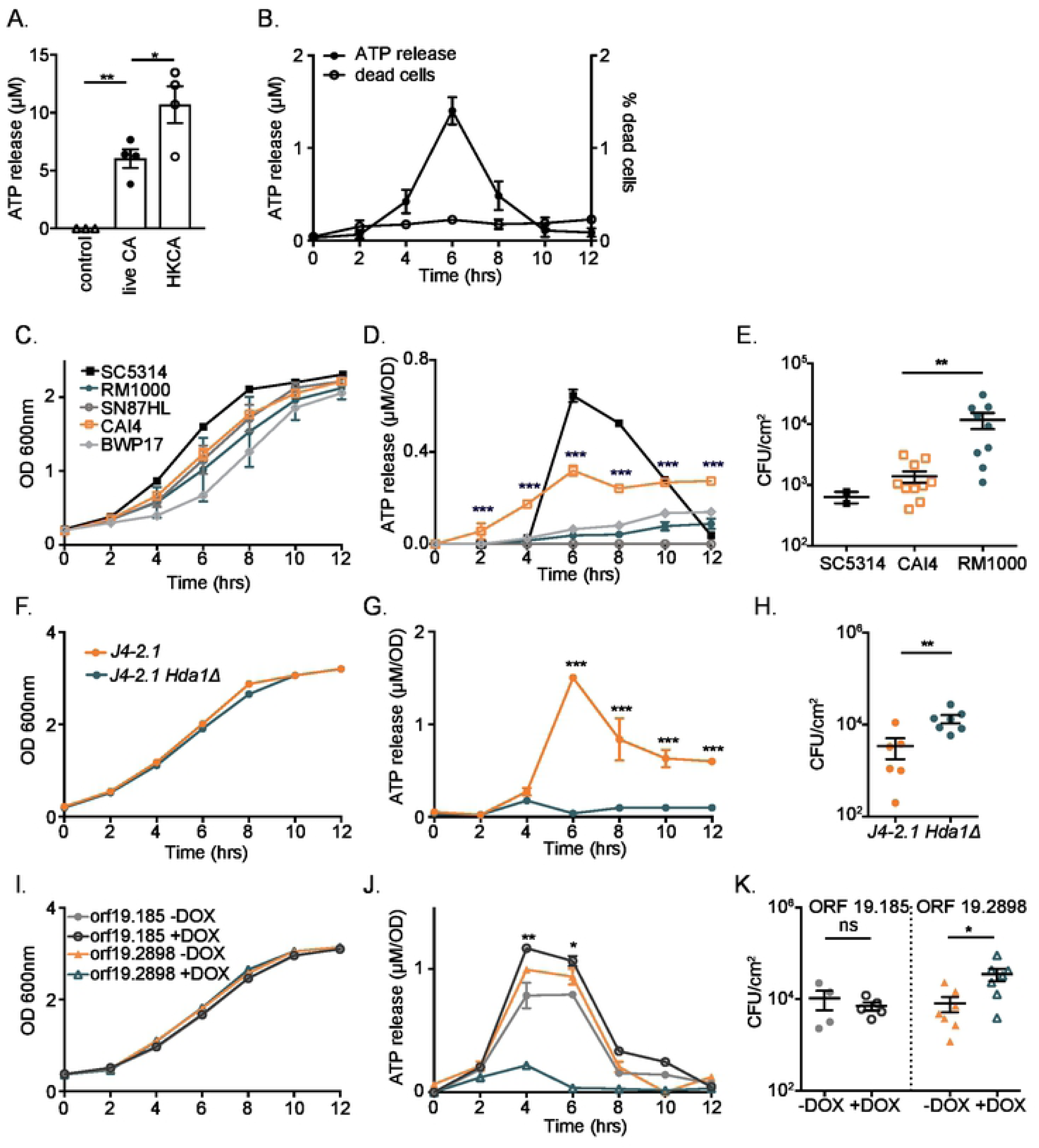
ATP is released from live *C. albicans* and correlates inversely with infectivity. **(A)** Concentrations of ATP were determined from supernatants of 4×10^8^ SC5314 *Candida* in PBS following either 60 minutes incubation at 0° (control), shaking incubation at 30° (live Ca) or incubation at 100° (HKCA). **(B)** Concentrations of ATP were determined from YPD broth during 30° shaking incubation of SC5314 at the indicated time point (black circles, left axis). The percent of dead yeast as determined by propidium iodide staining and flow cytometry at the indicated time point is shown (open circles, right axis). **(C)** Laboratory strains SC5314, RM1000, SN87HL, CAI4 and BWP17 were tested for growth at the indicated time point based on OD_600_ nm during 30° shaking incubation. **(D)** The concentration of ATP was determined from the same samples in (C) and is shown as the concentration of ATP normalized to OD_600_. **(E)** WT mice were epicutaneous infected with SC5314 (black circles), CAI4 (orange squares), or RM1000 (teal circles). The number of CFU obtained from homogenized skin harvested 3 days post infection is shown. **(F)** Kinetic analysis of OD_600_ and **(G)** concentration of ATP in YPD broth during 30° shaking incubation is shown for the parental J4-2.1 (orange) and *Hda1*Δ (teal) strains. **(H)** CFU 3 days post epicutaneous infection with J4-2.1 (orange) and *Hda1*Δ (teal) is shown. **(I)** Kinetic analysis of growth based on OD_600_ during 30° shaking incubation of Doxycycline-inducible overexpression lines *orf19.185* (circles) and *orf19.2898* (triangles) in the absence (gray, orange) or presence (black, teal) of doxycycline is shown. **(J)** The concentration of ATP from samples in (I) is shown. Release of ATP is significantly reduced throughout growth in doxycycline supplemented YPD for *orf19.2898 (teal triangles),* but not for *orf19.185 (black circles)*. **(K)** CFU obtained from skin 3 days following epicutaneous infection of the samples in (I,J) is shown. The doxycycline induced reduction in ATP release in ORF 19.2898 (teal triangles) is associated with higher infectivity. Significance was calculated using Mann-Whitney for all infection models (E,H,K), a Student’s t-test was used for ATP release (A,D,G,J). *p<0.05, \*\**p*<0.001, \*\*\**p*<0.0001. See also Fig S2.

To determine whether the SC5314 strain is unique in its secretion of ATP, we compared growth kinetics and eATP secretion in a collection of common prototrophic SC5314-derived laboratory strains. Notably, SC5314 and CAI4 released eATP while RM1000, SN87HL and BWP17 released very little eATP (**Fig 2C-D**). We next compared the capacity of CAI4 and RM1000, two strains with similar growth kinetics, to infect mouse skin using an epicutaneous skin infection model [13,26]. On day 3 following infection, CAI4, which secretes eATP, resulted in lower colony forming units (CFU) when compared to RM1000, a genetically related strain, that does not secrete eATP (**Fig 2E**). These data suggest that eATP enhanced *C. albicans* host defense though there are many genetic differences between CAI4 and RM1000 that could also potentially explain these results.

We next sought to identify a single gene mutant that results in reduced eATP. Our prediction was that these mutants should result in higher levels of infection *in vivo*. We focused exclusively on mutants with reduced eATP since the predicted increased levels of infection would be unlikely to result from the effects of genetic mutation that affect other aspects of *C. albicans* biology. A library screen of *S. cerevisiae* identified several genes required for release of eATP [27]. The majority of available *C. albicans* orthologous mutant strains, however, were on genetic background that did not release eATP (e.g. BWP17). We were able to identify one orthologous mutant, *hda1^Δ/Δ^*, that showed normal growth kinetics compared with its parental strain (J4-2.1) but did not release eATP (**Fig 2F-G**). As predicted, on day +3 following epicutaneous infection, CFUs of the *hda1^Δ/Δ^* mutant were significantly higher than those from control J4-2.1 cells (**Fig 2H**). Finally, we screened a series of dox-inducible OE strains based on our prediction from the data in Peters *et al*. (2016) that they would be affected in eATP secretion [28,29]We found two clones that secreted appreciable amounts of eATP and showed similar growth kinetics. Incubation with 50 μg/ml of doxycycline reduced eATP release from P_TET_-ORF19.2898 but not from P_TET_-O-ORF19.185 (**Fig 2I-J**). Both strains were grown in YPD augmented with doxycycline or vehicle and used to epicutaneously infect mice. On day +3 following epicutaneous infection only P_TET_-ORF19.2898 grown in the presence of doxycycline showed enhanced CFU (**Fig 2K**). From these data using laboratory strains of *C. albicans*, we conclude that reduced levels of released eATP resulted in enhanced epicutaneous skin infection *in vivo*.

### C. albicans skin infectivity negatively correlates with ATP secretion

To extend our findings beyond laboratory strains of *C. albicans*, we screened for release of eATP across 89 well characterized clinical isolates [30]. Isolates were categorized into 3 groups: commensal, if they were obtained from asymptomatic barrier sites (i.e. mouth, GI, or vagina); superficial if they were obtained from symptomatic barrier sites (e.g. oral thrush, urine); or invasive if they were obtained from blood. Isolates were grown in YPD for 6 hrs and analyzed for OD_600_ nm, levels of extracellular ATP in the YPD broth, and intracellular ATP levels from washed cells. Growth appeared relatively similar in most isolates (**Fig 3A**) but levels of eATP released into the broth varied across 3.5 logs and was independent of levels of intracellular ATP (**Fig 3B-C**). Notably, levels of eATP released by invasive blood isolates were approximately 10-fold lower than commensal and superficial isolates (**Fig 3B**). Release of eATP varied somewhat by genetic clusters (**Fig S3A**) suggesting that genetic differences determine levels of eATP. These data are consistent with our observations in laboratory strains that lower eATP is associated with greater infectivity *in vivo*.

**Fig 3:**
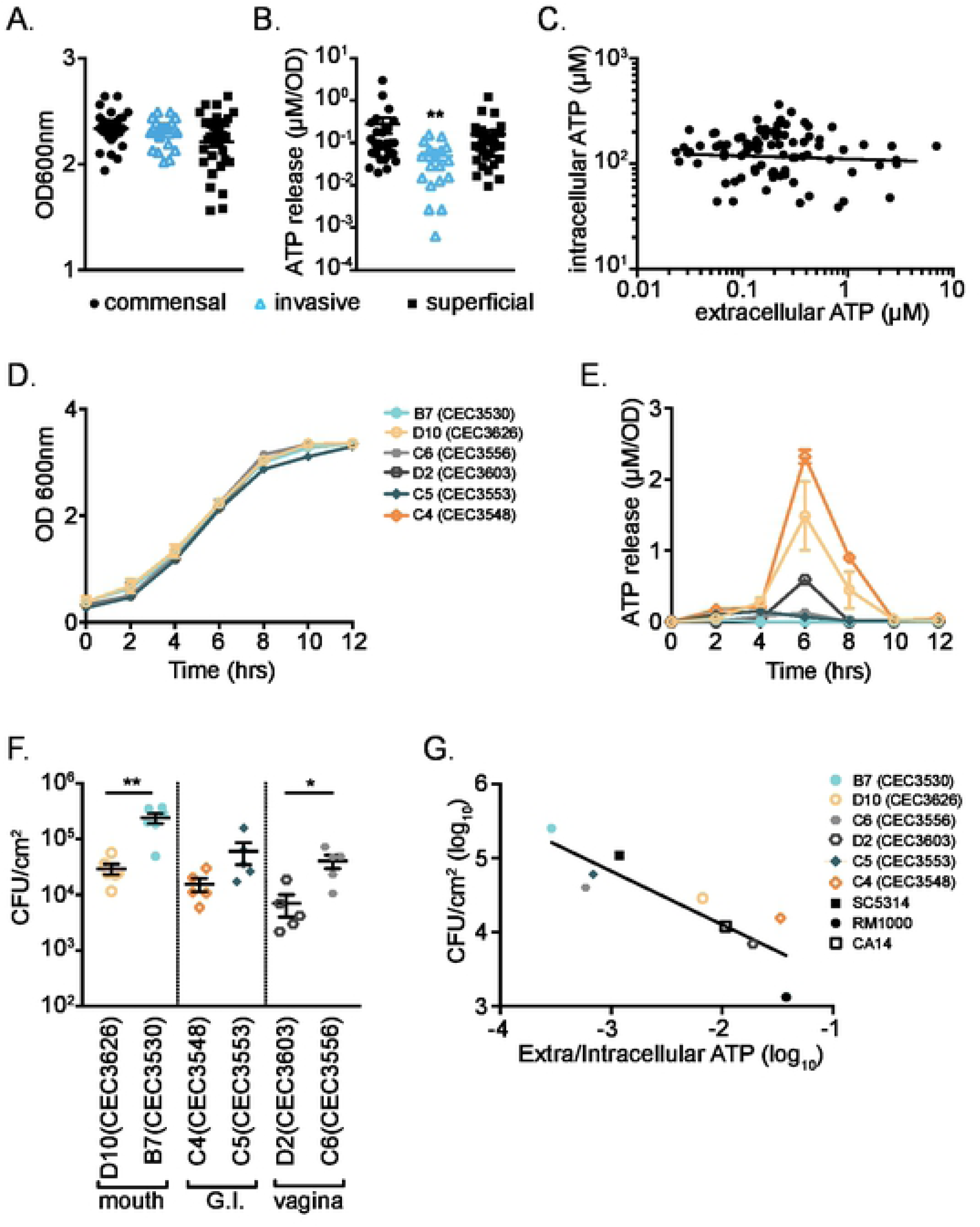
Clinical *C. albicans* isolates release ATP that inversely correlates with infectivity. **(A)** 89 clinical isolates, derived from superficial infections (black squares), blood infections (blue triangles), or commensal sites (black circles), were tested for growth by OD_600_ following 6 hrs 30° shaking incubation. **(B)** ATP released into the YPD broth of the same samples in (A) is shown. **(C)** Intracellular ATP concentrations from lysed *Candida* do not correlate with levels of extracellular ATP released from the same samples as in (A). **(D)** Representative commensal isolates from mouth (D10, B7), GI (C4, C5), and vagina (D2, C6) were examined for kinetic growth and **(E)** ATP release. **(F)** CFU on day 3 post epicutaneous infection with the indicated strain is shown. Isolates with high eATP (open symbols) have a lower CFU than strains with low eATP (closed symbols). **(F)** A strong inverse correlation between CFU following epicutaneous infection and the normalized concentration of eATP following 6 hours shaking 30° incubation is shown (p=0.002, Spearman r −0.8747). Each symbol represents data from an individual isolate (A-C), the average of 3 samples +/− SEM (D, E), or data from an individual animal (F). Significance was calculated using Kruskal-Wallace one-way analysis of variance (A) or Mann-Whitney (F). *p<0.05, \*\**p*<0.001, \*\*\**p*<0.0001. See also Fig S3.

To determine whether secretion of ATP correlates with skin infection, we selected paired high eATP- and low eATP commensal isolates from the mouth (D10:CEC3626, B7:CEC3530), GI tract (C4:CEC3548, C5:CEC3553) and vagina (D2:CEC3606, C6:CEC3556). These isolates transitioned normally from yeast to hypha when cultured in serum at 37° (**S3B Fig**), had similar growth kinetics (**Fig 3D**), but only C4, D10, and D2 released eATP (**Fig 3E**). Mice epicutaneously infected with high eATP strains from the mouth (D10), GI (C4), and vagina (D2) all showed lower CFU on day 3 following infection than their matched low eATP counterparts (**Fig 3F**). Moreover, the amount of eATP normalized to levels of intracellular ATP for every isolate tested has a very strong negative correlation with CFU (**Fig 3G**). Taken together, these data demonstrate that *C. albicans* releases eATP that varies greatly between strains. Moreover, low levels of eATP strongly correlate with increased infection efficiency suggesting that eATP released by *C. albicans* is recognized by the host and augments host defense.

### Recognition of C. albicans-derived ATP is mediated by P2RX2/3

Most immune cells in the skin express the purinergic receptors P2RX7 and/or P2RX4 (**Fig 4A**; reviewed in [31,32]). Of particular interest, is P2RX7, which is known to participate in the development of cutaneous Type-17 immune responses [33]. To determine whether host defense to *C. albicans* requires P2RX7, we epicutaneously infected wild type (WT) and *P2rx7*^−/−^ mice with a high ATP secreting strain (C4:CEC3548). On day 3 following infection, *C. albicans* CFU was equivalent in both groups indicating that P2RX7 is redundant for *C. albicans* host defense (**Fig 4B**). To test the requirement for P2RX4, WT mice were injected with 1mM 5-BDBD (a selective P2RX4 inhibitor) or vehicle intradermally (i.d.) starting on the day of *C. albicans* infection and twice daily continuing until harvest on day 3 (**Fig 4C**). Inhibiting P2RX4 did not affect *C. albicans* CFU.

**Fig 4:**
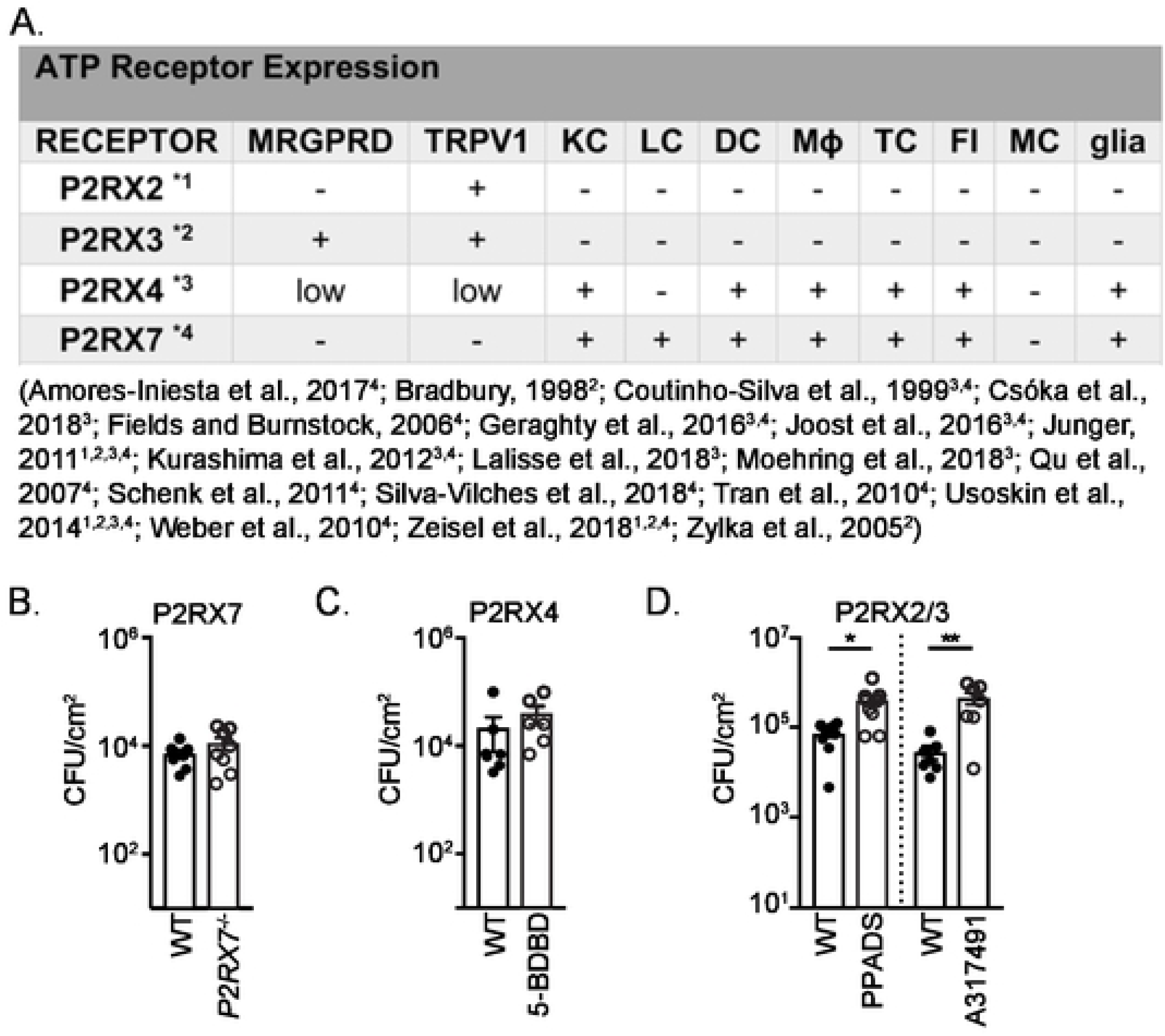
The ATP receptor P2RX2/3 is required for host defense to C. albicans. **(A)** Distribution of ligand gated ATP receptor expression across cell types found in skin (MRGPRD-expressing neurons, TRPV1-expressing neurons, keratinocytes (KC), Langerhans cells (LC), dendritic cells (DC), macrophages (MΦ), T cells (TC), fibroblasts (FI), mast cells (MC) and glia). References for ATP receptors are indicated by superscript in column 1: P2RX2 = 1, P2RX3 = 2, P2RX4 = 3, P2RX7 = 4. *1) [9,10,58] *2) [10,12,58] *3) [9,10,32,51,58–63] *4) [9,10,32,51,58,62–70]. **(B)** CFU isolated from skin 3 days following epicutaneous infection of WT or *P2Rx7*^−/−^ mice with the high ATP releasing clinical isolate C4 (CEC3548). **(C)** As in (B) except WT mice were infected and i.d. injected twice daily with 100μL vehicle or the P2RX4 antagonist 5-BDBD (1mM). **(D)** As in (C) except WT mice were i.d. injected twice daily with 100μL vehicle or the P2RX2/3 antagonists PPADS or A317491, both at 500µM. Each symbol represents data from an individual animal. Significance was calculated using Mann Whitney *p<0.05, \*\**p*<0.001, \*\*\**p*<0.0001.

In the skin, P2RX3 is primarily expressed by nonpeptidergic neurons particularly the subset identified based on expression of the Mas-related G-protein coupled receptor D (MRGPRD) [9,12,34]. To test the requirement for P2RX3, WT mice infected with *C. albicans* were injected i.d. throughout infection with two P2RX antagonists; the semi-selective agonist pyridoxal phosphate-6-azophenyl-20,40-disulphonic acid (P2RX1-3, P2RX5; PPADS), or the P2RX3 homomer- and P2RX2/3 heteromere-specific A-317491 [35,36]. Mice treated with either P2RX2/3 inhibitor showed significantly increased fungal burden compared with vehicle treated mice (**Fig 4D**). These data suggest that the cell types recognizing ATP during *C. albicans* infection are cutaneous sensory neurons. Based on our *in vitro* work (**Fig 1**) showing that eATP is not involved in release of CGRP, it is unlikely that CGRP-expressing TRPV1+ peptidergic neurons are the neuron subset responding to eATP. Rather, we hypothesize that MRGPRD+ nonpeptidergic neurons are the key cell type responding to *C. albicans*-derived eATP.

### Augmented host defense to C. albicans-derived ATP requires MRGPRD+ neurons

MRGPRD-expressing non-peptidergic neurons are a distinct subset of c-fibers that terminate within the epidermis and are characterized by binding IB4 and high expression of GFRa2, and P2RX3 [12,37,38]. We bred MRGPRD-Cre [39] with ROSA26.TdTomato (TdT) reporter mice to confirm appropriate expression of Cre in cutaneous neurons. As expected, TdT was highly expressed by cutaneous neurons terminating in the epidermis (**Fig 5A**). Immunofluorescent microscopy of DRGs also revealed that TdT was expressed primarily by cells binding IB4 and less by cells expressing CGRPα or TRPV1 (**Fig 5B and C**). We next examined Ca^2+^ flux following exposure to HKCA effluence in DRG cultured neurons from ROSA26.Tdt mice. Four distinct populations of cells could be identified (**Fig 5C**). The majority of effluence responding cells expressed TdT. Most did not respond to capsaicin indicating that these are MRGPRD+, TRPV1-nonpeptidergic neurons, but an overlapping capsaicin-responsive TdT+ population was evident. A small population of TdT-negative cells were capsaicin responsive indicating that these are likely TRPV1+ CGRP+ peptidergic neurons. A capsaicin-unresponsive TdT-population was also present.

**Fig 5:**
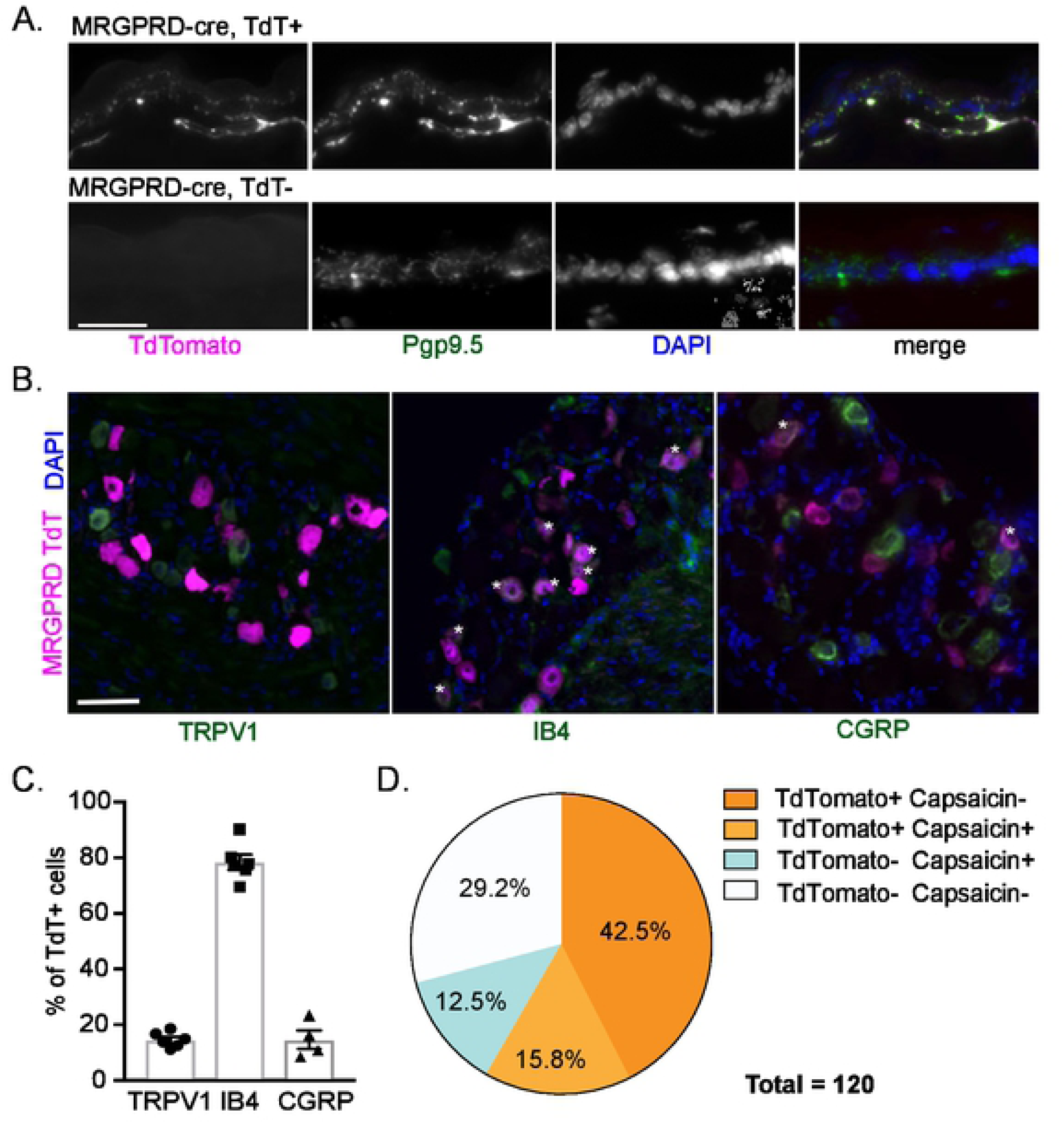
MRPGRD-expressing neurons respond to Candida. **(A)** Representative immunofluorescent microscopic images of unmanipulated skin stained with PGP9.5 and DAPI harvested from MRGPRD-TdT (top panels) or control ROSA26.TdT (bottom panels) mice is shown. MRGPRD-expressing neurons are evident throughout the epidermis. **(B)** Representative immunofluorescent microscopic images of unmanipulated DRG from MRGPRD-TdT mice labeled with anti-TRPV1 (left), IB4 (middle), or anti-CGRP (right). TdT expression (magenta) in lumbar DRGs overlaps (asterix) well with IB4, but less with TRPV1 or CGRP (green). **(C)** Summary of data in (B) showing the percentage of TdT+ cell bodies that express subset-specific markers. **(D)** The percentage of subtypes of DRG neurons (120 total) that flux Ca^2+^ in response to HKCA effluence is shown. DRG are divided based on expression of MRGPRD-TdT and response to capsaicin. Each symbol represents data from an individual animal. See also Fig S4.

To ablate MRGPRD neurons, we bred MRGPRD-Cre to ROSA26.DTR [39]. These mice allow for the inducible ablation of MRGPRD-expressing neurons following administration of *Diphtheria* toxin (DT). Immunofluorescent microscopy of epidermal whole mounts and transverse sections showed efficient ablation of epidermal GFRα2-expressing neurons (**Fig 6A, 6B**). We were able to confirm the absence of cre expression in tissues outside the nervous system, such as in the aorta (not shown) and intestinal tract **(S4A Fig)** as previously demonstrated with antibody labeling [40]. Analysis of DRGs revealed decreased numbers of GFRa2 and IB4 positive cell bodies consistent with a loss of nonpeptidergic neurons (**Fig 6A, 6C, S4B Fig).** We did not observe increase tail flick latency to heat, a characteristic behavioral screen for loss of TRPV1-expressing neurons **(S4C Fig).** We also did not observe alterations in numbers of immune cells in the skin or lymph nodes **(S4D,E, S5, and S6 Figs)**. Thus, MRGPRD-expressing neurons represent the dominant population of neurons responding by Ca^2+^ flux in response to HKCA effluence and they can be efficiently and selectively ablated in MRGPRD-DTR mice.

**Fig 6:**
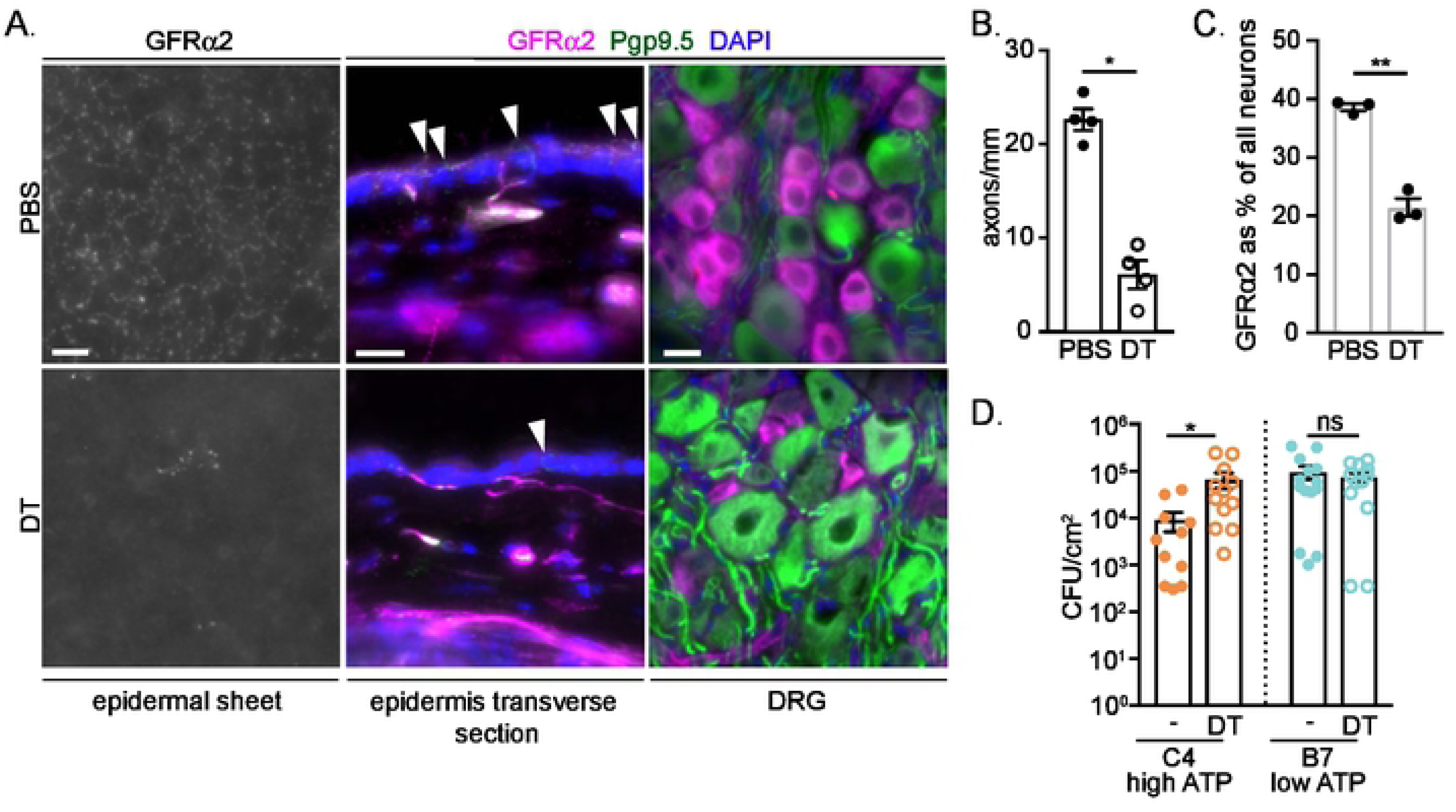
MRPGRD-expressing neurons are required for host defense against high ATP-secreting C. albicans. **(A)** MRGPRD-DTR mice were treated with DT to eliminate the subset of MRGPRD-expressing non-peptidergic neurons. The ablation of MRGPRD-expressing neurons is visualized by microscopic immunofluorescence using GFRα2 labeling in epidermal sheets of ear (scale bar 10μM), transverse sections of flank (scale bar 20μM), and in cross sections of the DRG (scale bar 20μm). All neurons are labeled with Pgp9.5 and nuclei with DAPI. Arrowheads indicate MRGPRD-expressing neurons in the epidermis. **(B)** Summary data from (A) showing the number of GFRα2 labeled neuron terminals crossing the dermal epidermal junction per linear mm is decreased after DT treatment. **(C)** Summary data from (A) showing the percentage of lumbar DRG neurons expressing GFRα2 is decreased by 44% after treatment with DT. (D) PBS (-) and DT treated MRGPRD-DTR mice were epicutaneously infected with high eATP (C4) and low eATP (B7) *C. albicans*. The CFU from skin isolated 3 days post infection is shown. Each symbol represents data from an individual animal. Statistical significance was determined by a Student’s t-test (B,C) and Mann-Whitney (D). Significance for all: *p<0.05, \*\**p*<0.001, \*\*\**p*<0.0001. See also Figs S4, S5 and S6.

To determine whether MRGPRD+ neurons participate in host defense against *C. albicans*, vehicle or DT treated MRGPRD-DTR mice were epicutaneously infected with either a high eATP strain (C4:CEC3548) or a low eATP strain (B7:CEC3530). On day 3 following infection, cutaneous *C. albicans* CFU was determined (**Fig 5G)**. DT treated mice in which MRGPRD+ neurons were ablated showed a significantly higher CFU compared with vehicle treated mice when infected with the high eATP strain. In contrast, DT and vehicle treated mice showed equivalent CFU when infected with the low ATP-secreting strain. Notably, CFU in DT treated mice infected with the high eATP strain was similar to mice infected with the low eATP strain. From these data we conclude that the augmented host defense that occurs as a result of *C. albicans*-derived eATP requires epidermal MRGPRD+ neurons. Moreover, the fact that CFU was independent of the presence of MRGPRD+ neurons during infection with a low eATP-secreting strain indicates that MRGPRD+ nerves do not participate in host defense in response to host-derived factors including keratinocyte-derived eATP or to *C. albicans*-derived factors other than eATP.

## Discussion

By initially examining the components of *C. albicans* responsible for activating cutaneous sensory afferent neurons, we found that neurons were not activated by yeast cell bodies but that a large number of non-peptidergic, TRPV1-negative neurons were activated by eATP. Some laboratory *C. albicans* strains released eATP and genetic alterations that reduced eATP secretion resulted in increased epicutaneous infectivity. Clinical isolates of *C. albicans* all released eATP with isolates from systemic infection showing greatly reduced levels of eATP. Infectivity across multiple isolates correlated well with increased eATP and decreased CFU recovered from infected tissue. Host recognition of *C. albicans*-derived eATP required P2RX3 which is expressed by epidermal non-peptidergic neurons. Ablation of MRGPRD non-peptidergic neurons resulted in greater infectivity for *C. albicans* that release eATP but had no effect on *C. albicans* releasing little, or no eATP. It is known that MRGPRD-expressing neurons respond to ATP [41] and these neurons have been implicated in modulating mechanosensation [42,43], histamine-independent itch [44] and extreme heat and cold sensation [45]. While these neurons are known to be involved in hyperalgesia following infection or injury [18,41], they have not previously been shown to play a role in the immune response to infection. Taken together, these data demonstrated that non-peptidergic cutaneous sensory afferents sense *C. albicans* through a novel, unappreciated mechanism that mediates host defense.

The release of eATP by *C. albicans* was unexpected. Although it is undetectable in some commonly used laboratory strains, all of the 89 clinical isolates we examined released amounts of eATP that were easily detectable. Furthermore, eATP levels were independent of levels of intracellular ATP, which is well known to be essential for growth, filament formation, and successful invasion [22]. ATP is released by *S. cerevisiae* and a mutational analysis revealed that many genes related to the secretory pathways were required for ATP efflux [27]. This suggests that release of eATP occurs through a defined mechanism and is not unique to *C. albicans*. Several bacterial species are also known to secrete eATP [46–49]. Notably, eATP from gut commensal bacteria is biologically active and is well documented to modulate the host immune response [19,20,50]. Interestingly, levels of eATP released by clinical isolates of *C. albicans* were highly variable. Invasive isolates identified in blood released less eATP and were over-represented in *C. albicans* clade 4 suggesting genetic control of eATP release. The observation that multiple microorganisms actively release ATP suggests that eATP may be beneficial to the yeast. In the case of *C. albicans,* we speculate that eATP may be related to a *C. albicans*-intrinsic process such as environmental sensing (e.g. the presence of host ATPases) or communication between individuals. In this scenario, strains of *C. albicans* at barrier surfaces that happen to have low eATP release are better able to avoid immune detection and are thus overrepresented in systemic infections.

The release of eATP by *C. albicans* augmented host defense as demonstrated with multiple genetically mutated laboratory strains where reduced eATP release did not affect growth or filamentation *in vitro* but did reduce infectivity *in vivo*. Similarly, the amount of eATP released from clinical isolates strongly correlated with reduced infectivity *in vivo* yet had no relationship with intracellular ATP levels or *in vitro* growth kinetics. Unexpectedly, recognition of eATP depended upon the P2RX2 or P2RX3 ATP receptors and was independent on P2RX4 and P2RX7. P2Rx7^−/−^ mice and WT mice treated with a P2RX4 antagonist, 5-BDBD, showed no change in *C. albicans* infectivity relative to control mice. Thus, there is no clear role for P2RX7 and P2RX4 in the recognition of eATP or for recognition of ATP from other potential sources in the host during *C. albicans* infection. In contrast, WT mice treated with two P2RX2/3-specific antagonists, PPADS and A-317491, had increased CFU in a skin infection model. In the skin, P2RX2/3 expression is largely restricted to neurons. P2RX3 is expressed in many non-peptidergic fibers [12] and very highly expressed in the MRGPRD subset [9]. Approximately 92% of MRGPRD-expressing neurons flux Ca^2+^ in response to 50µM ATP [41]. P2RX2/3 is not highly expressed in immune cells based on publicly available datasets (Immgen) or in other cells in the skin [51], nor is it highly expressed in glia [10]. We thus focused on the MRGPRD-expressing subset of nonpeptidergic sensory afferents.

In mouse, TRPV1-expressing and MRGPRD-expressing neurons are largely non-overlapping subsets. Only 5% of MRGPRD neurons respond to 1µM Capsaicin, and TRPV1 is expressed by only 9% of MRGPRD neurons [12,41]. We observed that 14.6% of DRG cells labeled in MRGPRD fate tracking mice co-express TRPV1. This raises the possibility that Cre driven by MRGPRD maybe transiently expressed on a small number of sensory afferent neurons during ontogeny. TRPV1 transcripts have been detected at low concentrations in MRGPRD neurons [52], which may also explain our observation of TRPV1 labeling in MRGPRD-cre driver expressing neurons. Despite this small amount of overlap, DT treated MRGPRD-DTR mice showed no attenuation in heat sensitivity. Thus, TRPV1 nociceptors remain functionally intact and MRGPRD-DTR mice efficiently and selectively ablate MRGPRD-expressing neurons in skin and in the DRG.

The ablation of MRGPRD-expressing neurons resulted in increased CFU of high eATP releasing, but not low eATP releasing, *C. albicans* strains. Thus, this neuron subset, which has dense free nerve endings in the epidermis, plays an important role in recognizing eATP and mediating the host response. This represents a novel function for this subset of sensory afferents. The absence of a phenotype with low eATP releasing *C. albicans* indicates that ATP from other potential sources during infection, such as keratinocytes, either does not activate these neurons or has no role in host defense. Possible mediators expressed by MRGPRD^+^ neurons that could modulate host defense are unclear, but a number of biologically active soluble mediators are expressed by these neurons [9,51].

In summary, we have identified that two major subsets of cutaneous sensory neurons recognize distinct *Candida*-associated PAMPS and together provide synergistic host defense. The mechanistic details of this synergy as well as the benefit to *C. albicans* derived from eATP release and its genetic control all remains as exciting future areas of exploration.

## Methods

### Animals

Mice used in this study are described in supplemental methods. Male and female mice between the ages of 7-13 weeks were used. Mice were housed in microisolator cages and fed irradiated food and acidified water.

### *Candida* strains and culturing

*C. albicans* SC5314, lab strains created from SC5314 [53], and lab strains created from clinical isolate SZ306, were all included in this study (see Supplemental Methods Table 1). Additional clinical isolates of different origins were previously described [30]. To measure growth, *C. albicans* was cultured in liquid YPD at 30°C/ 250 rpm overnight ∼16 hrs. These cultures were then diluted 1:50 in fresh YPD and incubated as above. Optical density was determined at the indicated time points. P_TET_-driven over-expression strains were cultured in YPD containing 50 μg/mL doxycycline and in the same medium without doxycycline as a control [28]. For induction of hyphae in liquid medium, *C. albicans* was sub-cultured for 3 hrs at 37°C in RPMI 1640 medium plus 10% (vol/vol) heat-inactivated FBS [54].

To stimulate neurons in culture, SC5314 was grown, as above, to ∼OD1.5, and washed three times in neuron recording buffer (NRB) to remove residual YPD. *Candida* was then resuspended in 10 mL NRB and heat-killed in a water bath for 30min-1hr at 100°C (assay dependent). HKCA was centrifuged at 3000RPM/5min and the NRB in which they were heat-killed – what we call effluence, was collected. HKCA were washed again and resuspended in NRB to a concentration of 1×10^7^/mL. HKCA and effluence were stored at −80°C until use. A similar method was used to create HKCA in PBS for ATP measurement.

### Neuron cultures

The caudal 6 thoracic DRGs and all 6 lumbar DRGs were dissected out and cultured for Ca^2+^ imaging, while all thoracic and lumbar DRGs were used for CGRP release assays. Dissection and methods for culture are based on those described in [55]. Following euthanasia and perfusion with ice cold Ca^2+^/Mg^2+^-free HBSS, isolated ganglia were placed into cold HBSS. In a two-step enzymatic digestion, DRGs were transferred to 3mL of filter-sterilized 60U papain/ 1mg L-cys/ HBSS buffered with NaHCO_3_ at 37°C for 10 minutes, then to filter sterilized collagenase II (4 mg/mL)/dispase II (4.67 mg/mL) in HBSS for 20 minutes at 37°C. Enzymes were neutralized in Advanced DMEM/F12 media with 10% FBS and 1% penicillin/streptomycin. Neurons were dissociated in media with trituration through a series of increasingly smaller-bored fire-polished glass pipettes. Cells for Ca^2+^ imaging were plated overnight in Advanced DMEM/F12 (as above) on poly-d-lysine coated 12mm round coverslips (Corning, Corning, NY). Cells for CGRP release assays were plated at a concentration of 20,000 cells/well in a 12 well plate in media supplemented with 10 ng/mL NGF 2.5S (Harlan, Indianapolis, IN) or 7S (ThermoFisher, Pittsburgh, PA) and cultured for ∼48 hr.

### Calcium imaging

Coverslips coated with neurons were transferred to a dish containing 2 mL 0.5% BSA in normal recording buffer (NRB: 1x HBSS with1.4mM NaCl, 0.9mM MgCl2, pH 7.4, osmolarity to 320 mOsm with sucrose), with 0.025% Pluronic F-127 (Promocell, Heidelberg, DE) and 1µM Fura-2AM (Molecular Probes, Eugene, OR), then incubated 30 minutes in the dark at 37°C, 5% CO_2_. Labelled neurons were placed in a 32°C heated recording chamber and superfused with NRB. Data was acquired on a PC with Metafluor software (Molecular Devices) via a CCD camera (RTE/CCD 1300; Roper Scientific). The ratio of fluorescence emission (510 nm) in response to 340/380 nm excitation (controlled by a λ 10-2 filter changer; Sutter Instrument) was acquired at 1 Hz during drug application. High K^+^ (50mM KCl), 1µM Capsaicin and 10µM ATP were applied using a computer-controlled perfusion fast-step system (switching time <20 ms; SF-77B perfusion system; Warner Instruments). Effluence, HKCA, and PPADS (10µM) were applied gently by hand using a 1mL insulin syringe with attached 0.28 mm diameter polyethylene tubing (Becton Dickinson, Franklin Lakes, NJ). At least 7 minutes was allowed between repeated stimulations with ATP or effluence. A 2s application of high K^+^ was used to evoke a Ca^2+^ transient. Cells unresponsive to high K^+^ were not analyzed.

To identify MRGPRD-expressing neurons MRGPRD-cre expressing mice were crossed to ROSA26.Tdt reporter mice. MRGPRD-expressing neurons were identified using the 538 laser, prior to excitation with 340/380 nm to determine which cells respond to effluence. An image of Tdt expressing cells was overlaid with the 340/380 image to determine what percentage of effluence-responsive neurons are MRGPRD neurons.

### CGRP release assay

The CGRP release assay was performed as previously described [13], but with slight modifications. After 48 hours in culture, 20,000 neurons/well were washed with 1xHBSS for 5 minutes and then incubated in NRB for 10 minutes to measure basal release. Treatments including 1×10^7^ SC5314 HKCA in NRB, effluence, 1U apyrase treated effluence, and 100µM αβ-me-ATP, were made for 10 minutes and the supernatant collected prior to lysing the cells in lysis buffer (20mM Tris, 100mM NaCl, 1mM EDTA, 0.5% Tx). CGRP in the supernatant was measured using the CGRP EIA kit (Cayman Chemical, Ann Arbor, MI) as per the manufacturer’s instructions using Gen5 1.10 software and a BioTek Epoch microplate Spectrophotometer (BioTek, Winooski, VT).

### Determination of ATP level of *C. albicans*

Aliquots of supernatant were collected from *C. albicans* cultures during growth and centrifuged at 3000 rpm for 5 mins at 4°C to remove yeast. ATP level in supernatant was determined using ATPlite luminescence assay system (PerkinElmer, Waltham, MA), as previously described [48]. To determine intracellular ATP level, the *C. albicans* pellet at each indicated time was washed in PBS and incubated for 30 min with ATPlite lysis buffer at room temperature. The extract was then spun down at 3000 rpm for 5 mins and the supernatant was transferred to a fresh tube for the ATP assay. Measurements were made with SOFTmax Pro for Lmax 1.1 software and a Lmax Microplate Luminometer (Molecular Devices, San Jose, CA).

### Apyrase digestion of ATP

To remove ATP, effluence was treated with MilliQ H_2_0 (control) or 1U/mL Apyrase in MilliQ H_2_0 for 1, 2, 4, and 8h at 30°C then heat-inactivated 20min at 65°C. ATP was measured using the ATPlite luminescence assay system (PerkinElmer, Waltham, MA). One hour treatments were determined to be sufficient for efficient ATP removal and thus were used in neuron stimulation assays.

### Skin Infection Model

Skin infection was performed as previously described [26]. Mice were anesthetized with ketamine and xylazine in sterile PBS (100/10 mg/kg body weight), their backs shaved with an electric clipper, and then chemically depilated with Neet sensitive skin hair remover cream (Reckitt Benckiser, Slough, England, UK). The stratum corneum was removed with 15-17 strokes using 220 grit sandpaper (3M, St Paul, MN). 2×10^8^ *C. albicans* in 50 μl of sterile PBS was then applied to the upper back.

For ATP receptor antagonist treatments, mice were injected intradermally across 5 sites on the back with a total of 100µL antagonist (500µM PPADS, 500µM A317491 in PBS or 1mM 5BDBD in DMSO/PBS) or PBS (with DMSO for 5-BDBD control) after sandpapering, but before application of *Candida.* Treatments were additionally given twice-daily at 12 hour intervals on d+1 and d+2. Skin was harvested 3-days post-infection, homogenized, and serially diluted onto YPAD plates. Plates were incubated at 30°C for 24-48 hours to determine colony forming units (CFU)/cm^2^.

### Immunofluorescent microscopy

For sectioned tissue, dissected mouse skin or DRGs were fixed in 4% paraformaldehyde (PFA) at RT for 1hr, washed twice in PBS, cryoprotected in 12% then 25% sucrose in PBS, and embedded in OCT at −80°C. DRGs and skin were cut at 14 μM and mounted onto Superfrost plus slides. For epidermal sheets, dorsal and ventral halves of the ear were separated and pressed epidermis side down onto double sided adhesive (3M, St. Paul MN). Slides were incubated in 10 mM EDTA in PBS for 45min, allowing physical removal of the dermis, before being washed 2x in PBS and fixed 30 minutes in 4% PFA in PBS before immunolabeling [56].

Primary antibodies included: 1:200 goat anti-GFRα2 (R&D Systems, Minneapolis, MI), 1:200-1:400 rabbit anti-PGP9.5 (Thermo, Pittsburgh, PA), 1:200 rabbit anti-TRPV1 (Alomone Labs, Jerusalem, Israel), 1:800 rabbit anti-CGRP (Sigma-Aldrich, St. Louis, MO). Pre-Conjugated antibodies and cell labels were used at 1:100-1:200 and include anti-MHCII AF488, anti-CD3 AF647, anti-GFP AF488 (Biolegend, SanDiego, CA), anti-tubulin BIII AF488 (eBioscience, San Diego, CA), and IsolectinB_4_-AlexaFluor 647 (Molecular Probes, Eugene, OR).

Secondary antibodies were used at 1:400 and include: goat anti-rabbit AF555, donkey anti-goat AF568, and donkey anti-rabbit AF647 (all Invitrogen, Carlsbad, CA). Antibodies were diluted in antibody diluting buffer (1% BSA, 0.1% Tween20 in PBS) with 5% normal goat or donkey serum. Tissues were mounted with Vectashield containing DAPI (Vector Laboratories, Burlingame, CA) and imaged using an Olympus fluorescent microscope (Olympus Corporation, Tokyo, Japan).

### Denervation

Resiniferatoxin (RTX; LC Laboratories, Woburn, MA) was injected on consecutive days at escalating doses of 30 mg/kg, 70 mg/kg and 100 mg/kg. Injections were made subcutaneously into the flank of four week old mice as previously described [57]. Four week old mice were given two intraperitoneal injections of 1 µg of Diptheriatoxin (DT; List Biological Laboratories, Campbell, CA) in 100µL PBS at three to four day intervals. Littermate control mice were treated on the same schedule with intraperitoneal injections of 100µl PBS.

### Cell/ Neuron Quantification

Quantification of cre-mediated eYFP+ co-expression with antibodies against TRPV1, IB4, and CGRP in TRPV1-TdT DRG neurons were each determined for 9 DRGs pooled from 2 mice. Data is represented as the percentage of cells expressing the indicated marker that co-express TdT. Quantification of neuron loss after DT treatment was characterized as a percentage of IB4 or GFRα2 neurons in β-tubulinIII labeled cell bodies. At least 5 sections from lumbar neurons of 3 DT treated and 3 PBS treated mice were analysed. Langerhans and DETC cells were counted from at least two separate images for each of 3 PBS and 3 DT treated mice. In cross sections of the ear, the number of GFRα2 labeled neuron terminals crossing the dermal epidermal border was pooled for 4-5 images from 4 mice for each treatment condition. ImageJ imaging software was used to quantify the cell number.

### Flow cytometry

Single cell suspensions were obtained from uninfected skin and lymph nodes as previously described [13]. Intracellular CD207 staining was performed using BD Bioscience Cytofix/Cytoperm kit (BD Biosciences, San Jose, CA). For extracellular staining, antibodies were diluted in staining media (3% Calf Serum, 5mM EDTA, 0.04% Azide) with Fb block. The following antibodies were used for skin: CD45.2 AF488, CD64 PE, TCRβ PE-Dazzle 594, CD3ε PE-Cy7, Gr1 AF 647, MHCII AF 700, CD11b BV421, TCRγδ BV510 (all Biolegend, SanDiego, CA), CD11c PerCPCy5.5 (Tonbo Biosciences, San Diego, CA), CD90.2 BUV395, CD8a BUV737 (both BD Biosciences, San Jose, CA). For lymph node staining we used: CD45.2 AF488, CD64 PE, Gr1 AF 647, MHCII AF 700, CD11b BV421, (all Biolegend, SanDiego, CA), CD11c PerCPCy5.5 (Tonbo Biosciences, San Diego, CA). Sample data was acquired on an LSRFortessa flow cytometer (Becton Dickinson, Franklin Lakes, NJ), and analysed using FlowJo software (TreeStar, Ashland, OR).

## Statistical Analysis

Statistical analyses were performed using GraphPad Prism software and is represented graphically as mean ± standard error. Chi squared analysis was used for comparisons of groups for Ca^2+^ imaging data, while Mann-Whitney and Student’s t test were used for comparisons between two groups. Kruskal-Wallace one-way analysis of variance (ANOVA) was used to compare corresponding groups as indicated.

### Ethics Statement

All animal experiments were approved by The University of Pittsburgh institutional care and use committee (IACUC #19126552). Mice were anesthetized with ketamine and xylazine in sterile PBS (100/10 mg/kg body weight). Euthanasia was performed using CO2.

## Acknowledgements

We thank the Division of Laboratory Animal Resources of the University of Pittsburgh for excellent animal care. This work benefitted from SPECIAL BD LSRFORTESSATM funded by NIH 1S10OD011925-01. This work was supported by the National Institute for Health (NIH) grants T32AI060525 (JAC), R01AR071720 (DHK, KMA, BMD), R01AR067187 (DHK) and European Research Council Advanced Award (340087 RAPLODAPT to J.B.).

## Author Contributions

T.N.E., S.Z., and D.H.K. designed and interpreted all experiments; A.L., J.A.C, S.M., P.Y.Z., performed experiments; B.H., J.B., M.B., A.R.M. and C.D. provided unique reagents, technical and conceptual assistance. H.R.K, S.L.G., K.M.A. and B.M.D. provided technical and conceptual assistance; T.N.E., S.Z. and D.H.K. wrote the manuscript and all authors edited it.

